# *A Toll pathway effector protects* Drosophila *specifically from distinct toxins secreted by a fungus or a bacterium*

**DOI:** 10.1101/2020.11.23.394809

**Authors:** Jianqiong Huang, Yanyan Lou, Jiyong Liu, Philippe Bulet, Renjie Jiao, Jules A. Hoffmann, Samuel Liégeois, Zi Li, Dominique Ferrandon

## Abstract

The *Drosophila* systemic immune response against many Gram-positive bacteria and fungi is mediated by the Toll pathway. How Toll-regulated effectors actually fulfill this role remains poorly understood as the known antimicrobial peptide (AMP) genes it controls are essentially active only against filamentous fungi and not against Gram-positive bacteria or yeasts. *BaramicinA* gene expression is transcriptionally regulated by the Toll pathway. *BaraA* encodes a polyprotein precursor that releases processed proteins into the hemolymph upon immune challenge. Here, we demonstrate that *BaraA* is required specifically in the host defense against *Enterococcus faecalis* and against the entomopathogenic fungus *Metarhizium robertsii*. It does so by protecting the fly from the action of distinct toxins secreted by Gram-positive and fungal pathogens but not by directly attacking them. Thus, in complement to the current paradigm, innate immunity can cope with toxins, effectively, through the secretion of peptides that are not AMPs, independently of xenobiotics detoxification pathways.

## Introduction

The study of host defense against infections has essentially focused on the immune response and the mechanisms used by the organism to directly attack, kill or neutralize invading pathogens. This dimension of host defense is known as resistance and in insects is mediated by antimicrobial peptides (AMPs) (1–4). However, there is a second complementary dimension known as disease tolerance or resilience whereby the organism is able to withstand and, in some cases, repair damages inflicted by the virulence factors of pathogens or the host’s own immune response (5–7).

Some instances of resilience have been reported in *Drosophila, e.g*., the removal of oxidized lipids by Malpighian tubules through the lipid-binding protein Materazzi or the requirement for *CrebA* in regulating secretion during the immune response (8, 9). One way to discriminate between resistance and resilience is to monitor the microbial burden of infected hosts. It will be increased during infection of immunodeficient as compared to immunocompetent hosts. In contrast, it will not change much in organisms with defective resilience, which will tend to succumb to a lower load of pathogens, as monitored by measuring the Pathogen Load Upon Death (PLUD) (10, 11).

In *Drosophila*, the Toll pathway is one of the two NF-κB pathways that regulate the systemic immune response to microbial infections and is required in the host defense against many Grampositive and fungal infections. It regulates the expression of more than 250 genes (9, 12–15). A few AMPs active against filamentous fungi have been identified (Drosomycin, Metchnikowin, Daisho) (16–18). However, effectors solely regulated by the Toll pathway able to attack pathogenic yeasts or Gram-positive bacteria *in vitro* have not been described so far. Mass-spectrometry analysis performed on the hemolymph of single immune-challenged flies has led to the identification of more than 30 peaks corresponding to Drosophila immune-induced molecules (DIMs) (19, 20). Some of them correspond to known AMPs whereas others belong to a family of 12 proteins that contain a domain known as the Bomanin domain (20, 21). Ten such *Bomanin* genes are located at the 55C locus and their deletion strikingly phenocopies the Toll mutant phenotype, being sensitive to filamentous fungi, pathogenic yeasts, and Gram-positive bacteria such as *Enterococcus faecalis* (21). There are some indications that short Bomanins that essentially contain only the Bomanin domain may be active against *Candida glabrata in vivo* (22).

Several DIMs are actually derived from a polyprotein precursor known as IMPPP and until recently their function was not understood. A recent study renamed this protein as BaramicinA (BaraA) and proposed that some of the derived peptides function as antifungal AMPs (23). Here, we report our analysis of *BaraA* mutants. While we confirm a sensitivity to entomopathogenic fungi, our data clearly establish a susceptibility also to *E. faecalis*, but not to other pathogens we have tested. Interestingly, the microbial burden does not appear to be altered in the mutants, from the beginning to the end of the infections. Our data indicate that the major function of *BaraA* is in the resilience against distinct toxins, a pore-forming toxin and a bacteriocin, respectively secreted by *M. robertsii* or *E. faecalis*.

## Results

The *BaraA* locus encodes a polyprotein precursor that is likely processed by a furin-like enzymatic activity, which leads to the release in the hemolymph of multiple DIM peptides. These peptides share extensive sequence similarity, except for the N-terminal DIM24 protein that defines an evolutionarily conserved independent domain (24) (Fig. 1A, B). Of note, *BaraA* lies next to the *CG18278* gene and the two genes are found as a perfect duplication in some wild and laboratory lines (24) (Fig. S1A, B).

**Figure 1.**
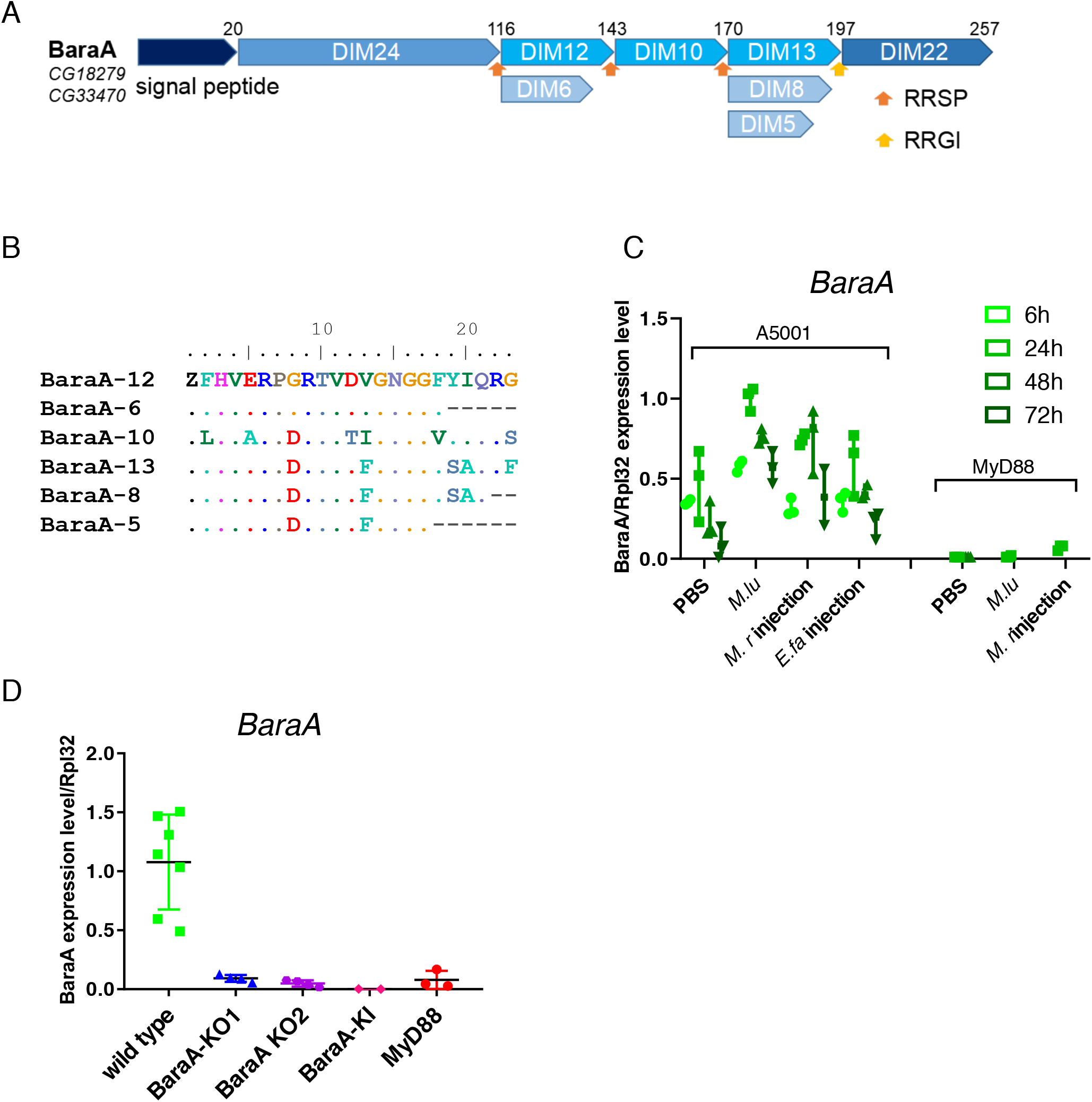
Structure of the BaraA precursor protein and induction of *BaraA* expression by an immune challenge. Schematic structure of the BaraA polyprotein. The name of the peptides derived from the processing of the precursor upon furin cleavage are shown as Drosophila-Induced Immune Molecules (DIM). The type of internal furin-like cleavage sites is indicated by orange and yellow arrows. **(B)** Alignment of the short DIM peptides derived from BaraA, referred to by their numbers. **(C)** Expression of the *BaraA* gene monitored by RTqPCR at various time points after the injection of the indicated microbes; *M. lu*: *M. luteus*; *M. r*: *M. robertsii*; *E. fa*: *E. faecalis*. **(D)** *BaraA* expression level measured by RTqPCR in wild-type, knock out (KO), knock in (KI), and *MyD88* flies, 24h after a *M. luteus* challenge.

*BaraA* gene expression is hardly detected in unchallenged flies in RNAseq studies (W. Wang, R. Xu, DF, unpublished data), in keeping with the absence of visible peaks after MALDI-TOF analysis on collected hemolymph from unchallenged flies (19). It is induced in a *MyD88*-dependent manner upon systemic infection by Gram-positive bacteria such as *Enterococcus faecalis* or by the fungus *Metarhizium robertsii* in two infection paradigms, injection of 50 conidia per fly or incubation of flies in a solution of spores at 5×10^4^/mL, the so-called “natural infection model” (25, 26) (Fig. 1C, Fig. S4C). In contrast, the *CG18278* gene does not appear to be induced by any of these challenges (Fig. S2A, B).

### BaraA *contributes to the host defense against* Enterococcus faecalis

In this work, we have generated two independent CRISPR-Cas9-mediated KO lines as well as a *mCherry* knock-in (KI) line (Fig. S1A,C, D). In these lines, the induction of *BaraA* expression by an immune challenge is hardly detected, both at the transcriptional (Fig. 1D) and protein level, even though some minor peaks can still be observed in the KO2 line (Fig. S3). We have also generated two *CG18278* KO lines, none of which revealed a sensitivity to an *E. faecalis* challenge, like one KD line (Fig. 2C-F). In contrast, we observed a significant susceptibility of *BaraA* mutant lines isogenized in the *w^A5001^* background after the injection of this Gram-positive bacterial strain (Fig. 2A). We did not measure any altered bacterial burden at any time in the KO and KI lines, even within 30mn after death (Bacterial Load Upon Death: BLUD (10); Fig. 2B-E).

**Figure 2.**
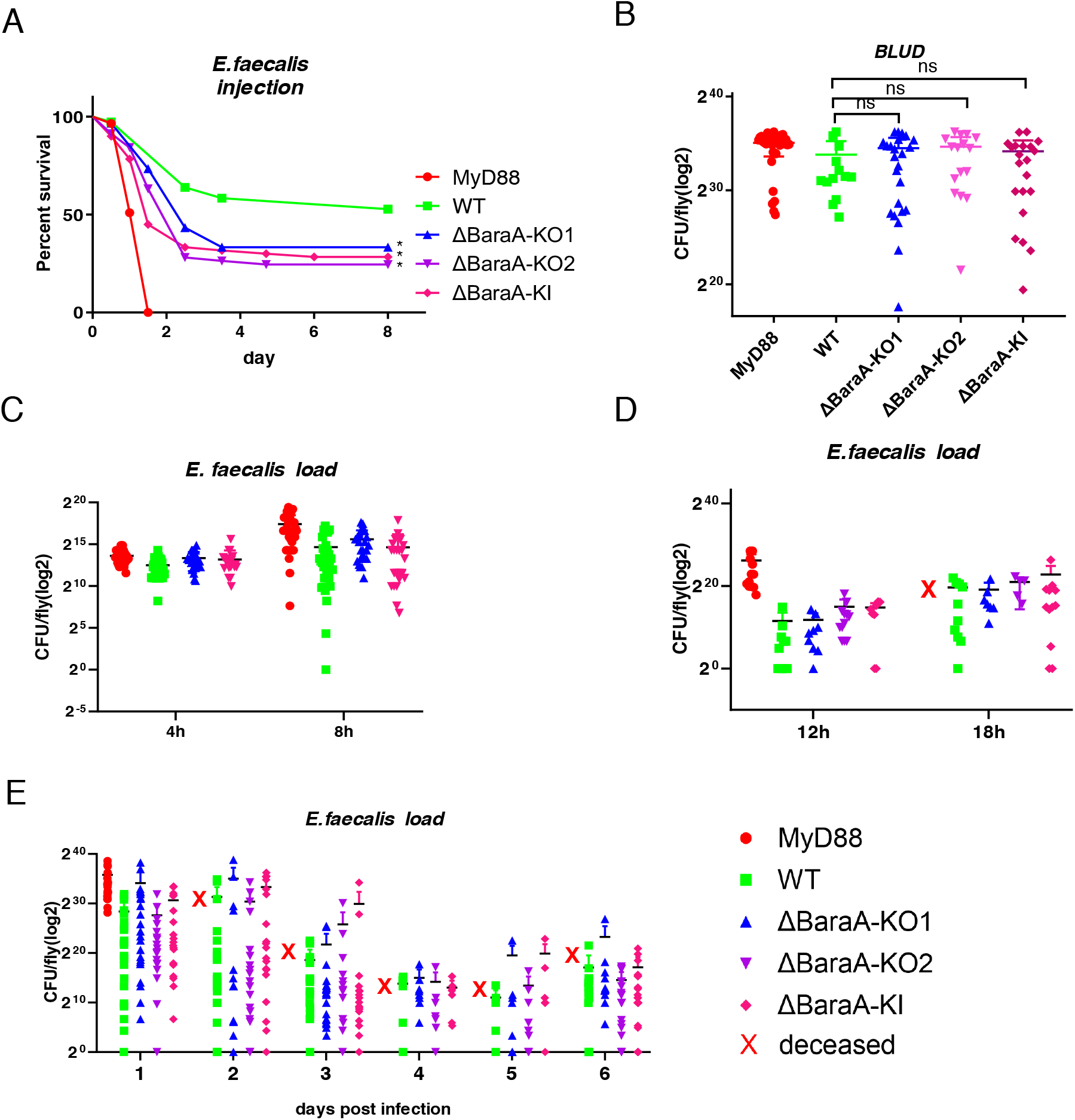
Susceptibility of *BaraA* mutant flies to *E. faecalis* infection. **(A)** Survival curves of the isogenic *BaraA* KO and KI flies infected with *E. faecalis;* one representative experiment out of five. The WT corresponds to a wild type *wA^5001^* line isogenized in parallel to the KO and KI lines, which behaves like the *wA^5001^* line used for isogenization. * p<0.05 **(B)** Bacterial load upon death (BLUD) of *E. faecalis* in WT and *BaraA* KO and KI lines. **(C-E)** Bacterial load of *E. faecalis* in WT and *BaraA* KO and KI lines from early time points (c, d) to days 1 to 6 after infection (e). No significant difference was detected between WT and mutants at each time point and BLUD. Captions apply to panels c-e. Pooled data from three independent experiments.

We conclude that the *BaraA* mutant lines display an intermediate sensitivity to *E. faecalis* infection and do not show a significantly altered bacterial burden.

### *The* BaraA *mutant is susceptible to* Metarhizium robertsii *infection only in the septic injury model*

The *BaraA* KO and KI lines consistently exhibited an intermediate sensitivity to the injection of 50 *M. robertsii* conidia (Fig. 3A). As for *E. faecalis*, we did not measure an increased microbial titer in these mutants during the infection; the Fungal Loads Upon Death (FLUD) were also similar (Fig. 3B,C). Interestingly, no susceptibility to *M. robertsii* in the natural infection model was observed, even though the *BaraA* gene is induced by this challenge (Fig. S4C, C’). We have also tested a panel of other bacterial and fungal strains and did not observe any sensitivity to those infections (Fig. S4).

**Figure 3.**
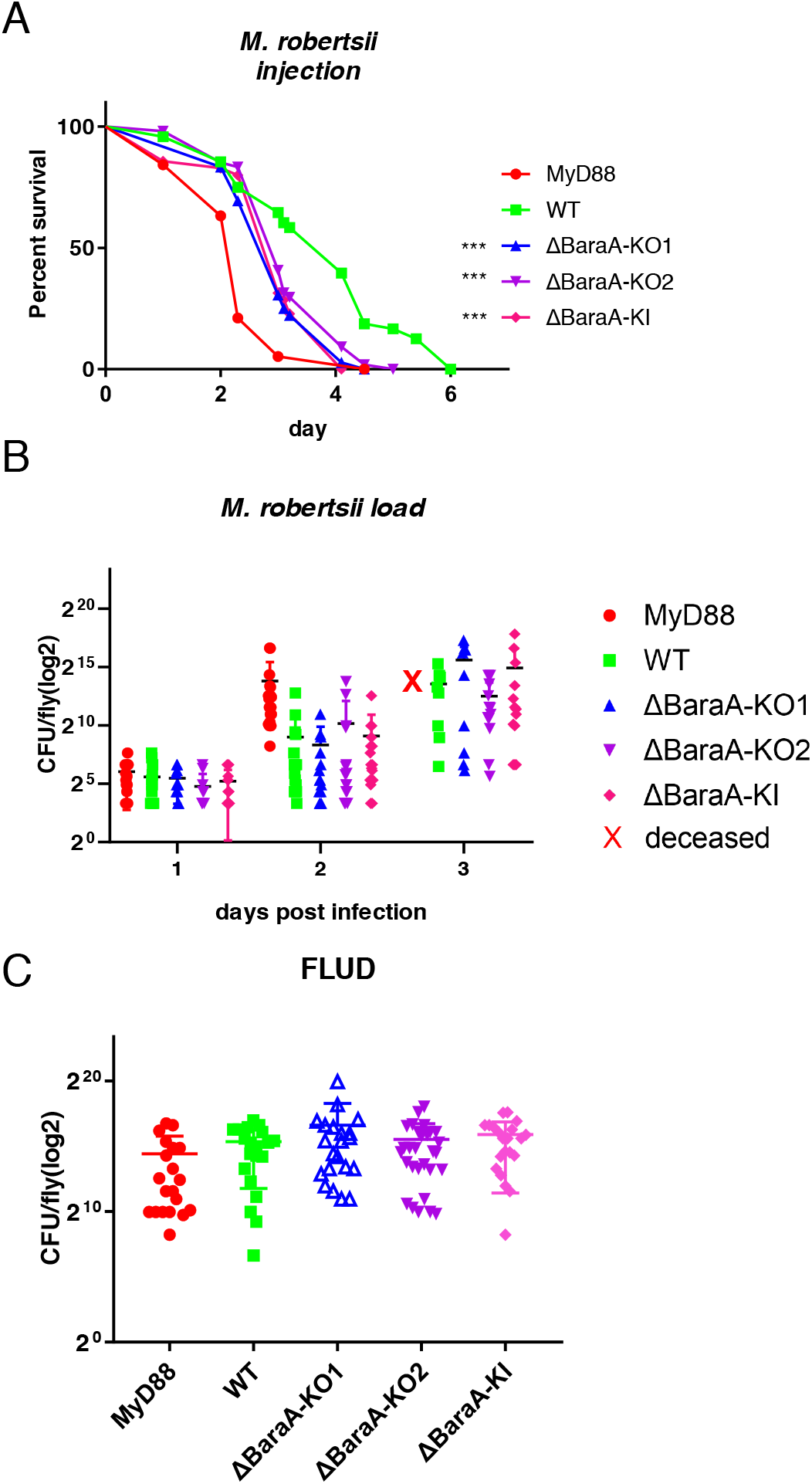
Susceptibility of *BaraA* mutants to *M. robertsii* infection. **(A)** Survival of isogenic *BaraA* mutants injected with 50 *M. robertsii* conidia. KO1, KO2 and KI performance were consistent across multiple independent experiments (n>3). Statistical significance between WT and the KO or KI mutants: *** p<0.001. **(B)** Kinetics over three days of the fungal load of the *BaraA* KO and KI mutants injected with 50 *M. robertsii* conidia. No significant difference was detected between WT and mutants at each time point. **(C)** Fungal load upon death (FLUD) of single isogenic *BaraA* KO and KI flies injected with 50 *M. robertsii* conidia. The fungal burden was measured within 30 minutes following the demise of the single flies. No significant difference between WT and the isogenic mutant flies was detected. Three independent experiments have been performed and pooled (B, C).

In conclusion, we have found that *BaraA* appears to be required rather specifically in the host defense against a bacterial opportunistic pathogen, *E. faecalis*, and an entomopathogenic fungus, *M. robertsii*. Interestingly, the microbial burden was not altered in the *BaraA* mutants for both infections, which indicates that *BaraA* is not required in the resistance against these pathogens.

### An overexpression strategy fails to confer an enhanced protection against several pathogenic bacteria or fungi

A complementary strategy to the loss-of-function analysis reported above consists in overexpressing the *BaraA* gene in a wild-type context thus determining whether it might constitute a limiting factor in host defense against infections. The overexpression of *BaraA* or of the sequence coding only for BaraA-encoded proteins, DIM22 excepted, using transgenic lines failed to enhance host protection against our panel of pathogens (Fig. S5A-C).

We next tested the overexpression lines in a sensitized *MyD88* background. As shown in the lower panel of Fig. S5A, *MyD88* is not required for the processing of the precursor into the DIM10, DIM12, DIM13 or DIM24 proteins by a putative furin. However, the carboxypeptidase activity needed to trim the DIM12 and DIM13 peptides into respectively DIM6 and DIM5/DIM8 appears to be dependent on Toll pathway activation since we did not observe these shorter peptides. We also conclude that the Bombardier activity needed to stabilize the expression of short Bomanins is not required for the stability of DIM10, DIM12, and DIM13 (27). Of note, the overexpression of C-terminal HA-tagged BaraA did rescue the *BaraA* sensitivity phenotype to both *E*. *faecalis* and *M. robertsii* when the transgene was expressed only at the adult stage (Fig. S5D-E), thereby demonstrating that the transgene is functional. In contrast, we failed to observe any rescue of the sensitivity of *MyD88* flies to a panel of pathogens by overexpressing *BaraA* or its derived peptide coding sequences in this mutant background (Fig. S5C). We conclude that the genetic overexpression of *BaraA* is not sufficient to confer additional protection against *E. faecalis* or *M. robertsii* in the context of a wild-type or Toll-deficient immune response.

### BaraA does not modulate the induction of the Toll pathway

Besides a potential role of effectors, proteins that are induced by immune signaling pathways may play a role in feed-back regulation of the signaling pathway. We therefore monitored Toll pathway activation using the steady-state mRNA levels of AMP genes known to be regulated by the Toll pathway such as *Drosomycin, Metchnikowin*, and *IM1* (28–30). As shown in Fig. S6, we did not observe any influence of the isogenized *BaraA* KO or KI null mutations over 48 hours.

### BaraA *protects* Drosophila *from the action of secreted microbial toxins*

Our data thus far are not compatible with a function for *BaraA* in resistance against *E. faecalis* or *M. robertsii*. The PLUD data are not indicative of a function in resilience. Indeed, we have excluded a potential role for *BaraA* in Malpighian tubule function and in the handling of metabolic stores (Fig. S7).

The concept of pathogen load and PLUD relies on the assumption that the virulence of the pathogen correlates with the microbial burden. We have recently established that the function of the Toll pathway in the host defense against *Aspergillus fumigatus* is not to directly fight off this pathogen, as immunodeficient flies are killed by a limited number of pathogens that are trapped at the injection site. Rather, we have discovered that Toll function in the host defense against *A. fumigatus* is to limit or counteract some of its secreted mycotoxins (see companion article). As mycotoxins, namely Destruxins, are also key virulence factors from *Metarhizium* entomopathogenic fungi (31), we therefore injected Destruxin A into wild-type and *BaraA* flies. Interestingly, *BaraA* KO and KI mutant as well as *MyD88* flies succumbed to a larger extent than wild-type flies to the injection of Destruxin A (Fig. 4A), a result confirmed in axenic flies (Fig. S8A). We next determined that *BaraA* mutants are not more susceptible than wild-type flies to a challenge with a *Dtx* mutant *M. robertsii* strain (32) (Fig. 4B). Taken together, these results suggest that a major function of *BaraA* in the host defense against *M. robertsii* is to alleviate or counteract the effects of Destruxins secreted by the fungus in the septic injury model.

**Figure 4.**
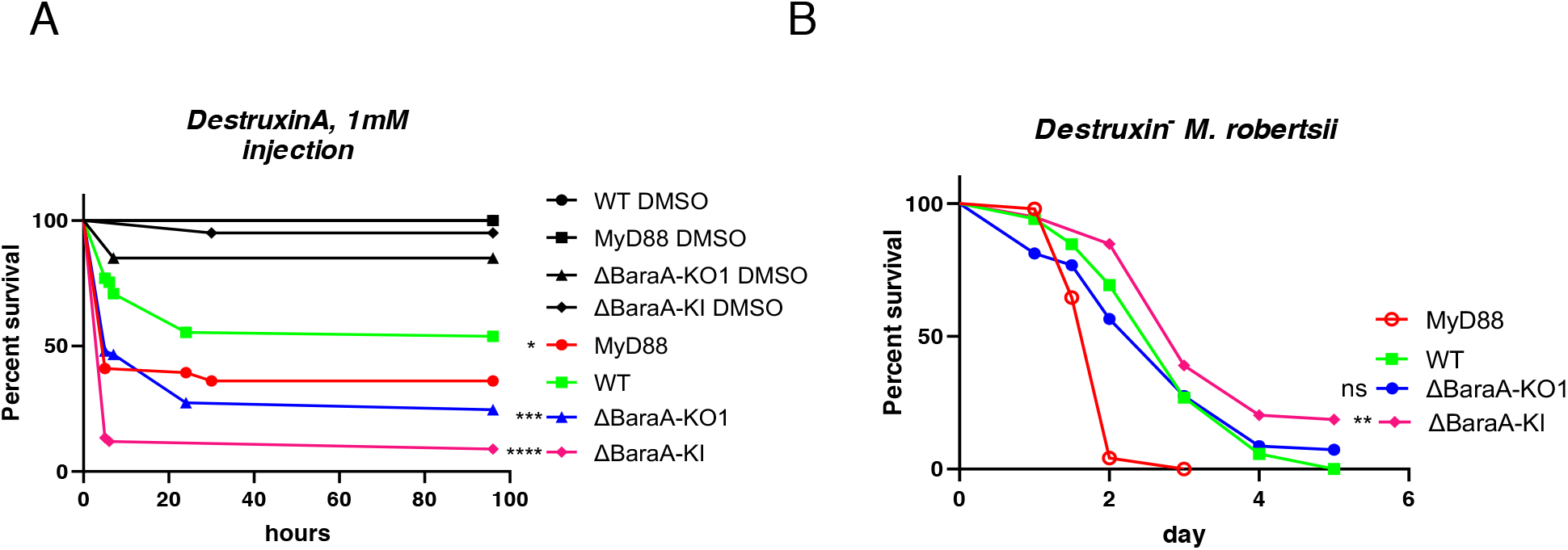
*BaraA*-dependent protection of *Drosophila* flies from the noxious effects of microbial toxins. **(A)** Mutant flies have been injected with 23nl, 1mM Destruxin A toxin. 20% DMSO was injected as vehicle control. One representative experiment out of more than three experiments. Statistically significant difference between wild type and *BaraA* mutants, * p<0.05, *** p<0.001, **** p<0.0001. **(B)** *BaraA* mutants were injected with 50 spores of *DestruxinS1^-^ M. robertsii* mutant strain in which Destruxins biosynthesis is blocked. No significant difference was observed between wild type and *BaraA* KO1 in more than three independent experiments. *BaraA* KI showed increased protection with respect to wild type in two out of five experiments, including this one; **p<0.01.

We then wondered whether *BaraA* might function in a similar manner in the host defense against *E. faecalis*. We therefore injected the *E. faecalis* culture supernatant into flies. Strikingly, whereas wild-type flies survived this challenge well, about 50% of *BaraA* and *MyD88* flies succumbed to this challenge (Fig. 5A). Even though the noxious activity in the *E. faecalis* supernatant was heat-resistant, it was nevertheless susceptible to proteinase K treatment, suggesting a protein component (Fig. S8B, C). Filtration experiments allowed us to determine that the toxic component of the supernatant can be recovered in a three to ten kD fraction (Fig. 5B). Interestingly, it has been reported that the bacteriocin enterocin O16 is an *E. faecalis* virulence factor in *Drosophila* (33). Enterocin O16 is also known as Enterocin V (EntV), which is heat-resistant and able to kill some lactobacilli strain as well as to inhibit the hyphal growth, the virulence, and biofilm formation of *C. albicans* (34, 35). It derives from the open reading frame found in the *ef1097/entV* gene, which encodes a preproprotein. This precursor protein gets cleaved by the GelE protease into a 7.2 kDa active peptide (35, 36).

**Figure 5.**
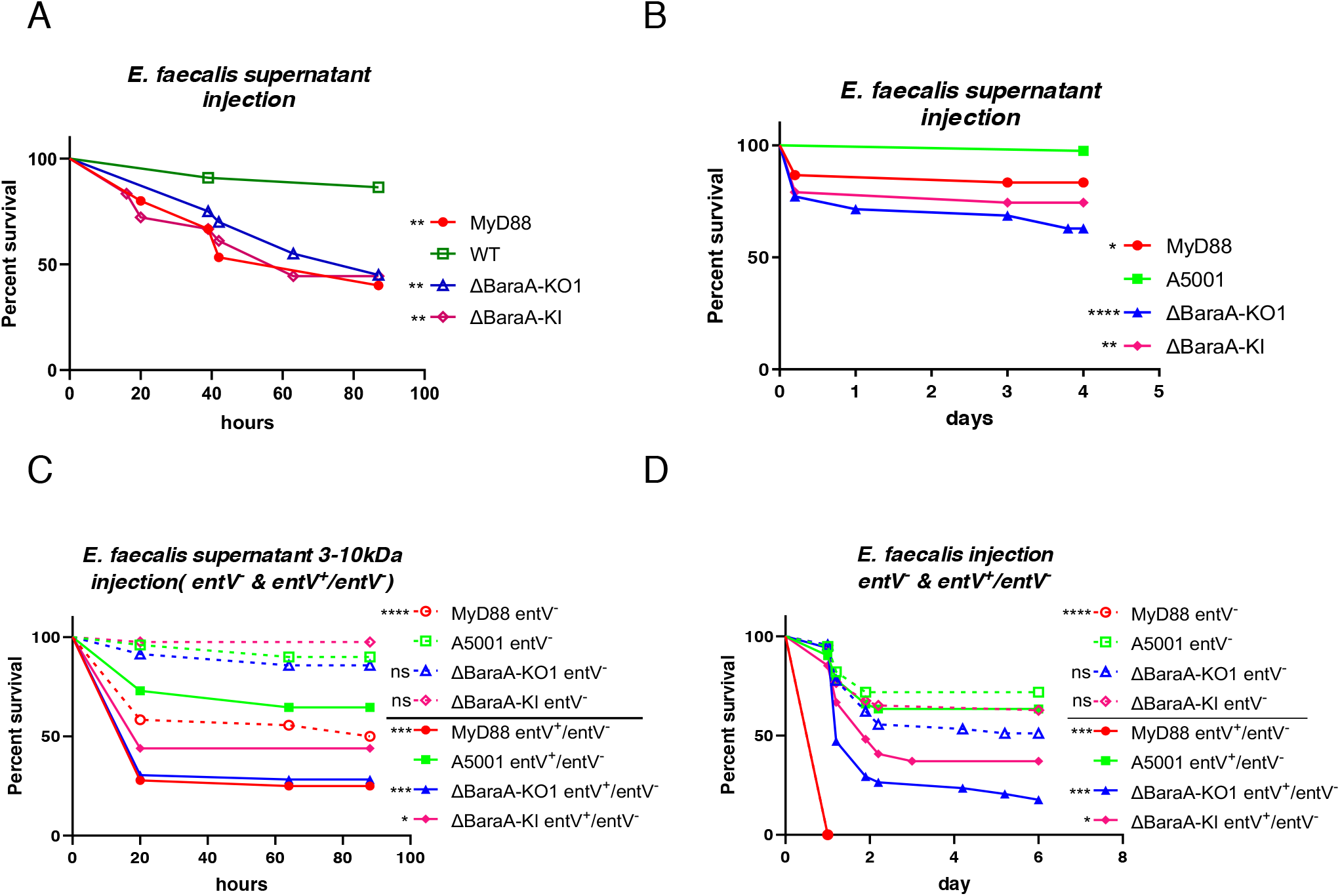
**(A)** Wild-type, *MyD88*, and *BaraA* KO1 and KI mutant flies were injected with the concentrated supernatant from overnight *E. faecalis* cultures. About 50% of mutant but not wildtype flies succumbed to this challenge; ** p<0,01. **(B)** Same as (A), except that the supernatant was size filtered to retain molecules ranging from three to ten kDa; * p<0.05, ** p<0.01, **** p<0.0001. **(C)** The 3-10kDa fraction supernatant from *E. faecalis* was collected from *entV*^-^ and *entV*^+^/*entV*^-^ strains. The supernatant from the *entV*^-^ strain killed *BaraA* mutants at the same rate as wild type whereas *BaraA* KO1 mutant flies were killed by the *entV^+^*/*entV*^-^ supernatant significantly faster than wild-type flies. For each condition (above or below the line in the caption), mutant flies are compared to wild-type flies submitted to the same challenge for statistical analysis. * p<0.05; *** p<0.001; **** p< 0.0001. **(D)** 0.5 OD, 4.6nl of the mutant and rescued *E. faecalis* strains were injected. Rescued strain *entV^+^/entV^-^* strain killed *BaraA* KO1 faster than wild-type flies, while no significant difference between wild type and *BaraA* mutant flies was detected upon *entV*^-^ infection. *BaraA* KI yielded the same results as *BaraA* KO1. For each condition (above or below the line in the caption), mutant flies are compared to wild-type flies for statistical analysis. * p<0.05; *** p<0.001; **** p< 0.0001. All experiments have been performed independently more than three times.

We therefore asked whether the toxic activity in the supernatant is still present when using a bacterial *entV*^-^ strain. We observed that the supernatant from the complemented *E. faecalis* strain *entV+/entV−* behaved as that from the wild-type bacterial strain, that is, it killed *MyD88* and *BaraA* KO and KI mutants more than wild-type flies. Strikingly, the supernatant from an *entV− E. faecalis* mutant strain no longer killed wild-type or *BaraA* mutants whereas *MyD88* flies displayed an intermediate sensitivity in this experiment (Fig. 5C). Conversely, the direct infection with the *entV*^-^ mutant *E. faecalis* strain did not differentially kill *BaraA* and *MyD88* mutants as compared to wildtype flies submitted to the same challenge (Fig. 5D). As expected, the complemented *E. faecalis* strain *entV+/entV−* behaved as the wild-type bacterial strain and killed the immuno-deficient *MyD88* and *BaraA* flies more than the wild-type control flies. In addition, a *gelE E. faecalis* mutant strain also did not kill *BaraA* flies faster than wild-type flies (Fig. S8D). We conclude that *BaraA* protects the flies from the action of the EntV bacteriocin.

Taken together, our data suggest that the major function of *BaraA* in *Drosophila* host defense is to protect the fly from specific secreted microbial toxins, whether of prokaryotic or eukaryotic origin.

## Discussion

Our analysis of the *BaraA* mutant phenotype revealed a susceptibility to specific pathogens and not to broad categories of microorganisms as is the case for Toll pathway mutants. Interestingly, we observed a susceptibility to *E. faecalis* and to *M. robertsii*, respectively a Gram-positive bacterium and an entomopathogenic fungus. For both pathogens, specific secreted virulence factors killed *BaraA* mutants whereas the phenotype of enhanced sensitivity to infection was lost when the corresponding virulence factor genes were mutated in the pathogen. Taken together, these results indicate that the major function of *BaraA* in *Drosophila* host defense is to protect it from the action of specific secreted toxins. Indeed, whereas in a companion article we showed that Toll pathway mutant flies are sensitive to *Aspergillus fumigatus* verruculogen and restrictocin, *BaraA* mutants did not exhibit any enhanced sensitivity to these mycotoxins.

A recently published study proposed that BaraA is involved in resistance to infection to entomopathogenic fungi as an AMP since, besides being sensitive to *Beauveria bassiana* and *Metarhizium rileyi*, they exhibit an increased *B. bassiana* load 48 hours after infection (23). In addition, BaraA-derived IM10-like peptides synergize with a membrane-active antifungal compound to kill *Candida albicans in vitro* (23). The fact that BaraA is a polyprotein that produces multiple DIM10-like peptides and that the *BaraA* locus is found to be duplicated in about 14% of wild-type *Drosophila* strains caught at one location is in keeping with this possibility. Because BaraA encodes a polyprotein precursor, we cannot formally exclude such an AMP function for one or several of these BaraA derivatives. Indeed, whereas we have shown a function for some specific 55C *Bomanins* in the resilience to *A. fumigatus* mycotoxins, it is known that at least some *Bomanin* genes are required for resistance to *E. faecalis*, a finding we have directly confirmed for at least one 55C *Bomanin* gene (37). We note that if DIM10-like peptides indeed act as AMPs, they would need to be specific for microbial strains as documented in this study. Very specific antibacterial functions for some *Drosophila* AMPs have been documented (38, 39); however, we are not aware of AMPs having dual specificities against both particular bacterial and fungal species. In our view, the finding of a loss-of-virulence of bacterial and fungal toxin mutants in *BaraA* mutant flies supports the concept that BaraA’s major function is to neutralize or counteract the action of specific secreted microbial toxins in the case of *E. faecalis* or *M. robertsii* infections, in as much as we did not detect an enhanced microbial load of these pathogens in *BaraA* mutants.

A study on the evolution of BaraA as well as two related genes suggests that the core domains of these three proteins is the N-terminal DIM24 domain, which is associated with only two DIM10-like domains in BaraB and none in BaraC (24). The expression of *BaraB* has been reported not to be induced by an immune challenge, a finding we have independently confirmed. We did not find a susceptibility of *BaraB* KO mutant to *E. faecalis* infection (unpublished data). Thus, it is likely that the DIM24 domain may have a function distinct from the DIM10-like peptides. Unfortunately, our overexpression experiments in wild-type or *MyD88* mutant backgrounds failed to reveal any protective phenotypes, thus precluding us from drawing any conclusions on the role of specific BaraA-derived peptides. We can only speculate that DIM24 might be involved in counteracting the action of microbial toxins whereas DIM10-like peptides might act as AMPs or opsonins.

The exact mode of action of BaraA in the resilience to microbial toxins remains to be characterized, in as much as it acts against distinct types of toxins. Destruxins have been isolated 60 years ago and they appear to act as ionophores that deplete cellular ions such as H^+^, Na^+^, and K^+^ through the formation of pores in the membranes in a reversible process (40). Bacteriocins also form pores on the membrane of targeted bacteria but the mechanism of action of EntV on eukaryotic cells remains unknown. Thus, it is unclear whether BaraA would act directly on both toxins, counteract a common process triggered by Destruxins and EntV such as intracellular ion depletion, or indirectly alters the physiology of cells exposed to the action of these toxins. It will be important to determine how the toxins act on the host and whether they target preferentially some tissues.

A specificity of the Toll pathway is that it is required in the host defense against both prokaryotic and eukaryotic pathogens. As compared to the IMD pathway, one interesting feature is that the Toll pathway can be activated by proteases secreted by invading pathogens (41–43). It is interesting to note here that the function of BaraA against two distinct secreted virulence factors, likely pore-forming toxins, provides another point of convergence for the dual role of the Toll pathway, this time at the effector level. It is thus an open possibility that one of the selective pressures that shaped the function of the Toll pathway would be the need to cope with pathogens secreting virulence factors in the extracellular compartment.

Taken together with the companion article, our work underscores that the Toll pathway mediates resilience against the action of multiple toxin types such as pore-forming toxins, ribotoxins or tremorgenic toxins, which are mediated by specific Bomanins or BaraA-derived proteins. It is likely that other uncharacterized effectors are able to counteract other toxins to which *Drosophila* flies are exposed to in the wild. In contrast to the current paradigm according to which secreted peptides act as AMPs, our discoveries illustrate a novel concept, the ability of the innate immune system to counteract secreted microbial virulence factors. Future studies should determine whether this capacity has also been selected for throughout evolution.

## Materials and Methods

### Fly strains

Fly lines were raised on media at 25°C with 65% humidity. For 25 L of fly food medium, 1.2 kg cornmeal (Priméal), 1.2 kg glucose (Tereos Syral), 1.5 kg yeast (Bio Springer), 90 g nipagin (VWR Chemicals) were diluted into 350 mL ethanol (Sigma-Aldrich), 120 g agar-agar (Sobigel) and water qsp were used.

*w^A5001^* (44) and *yw* flies were used as wild type controls. The positive controls for infection assays for Gram-positive/fungal infections and Gram-negative infections were respectively *MyD88* and *key* in the *w^A5001^* background. Where stated, mutant flies were isogenized in the *w^A5001^* background. For RNAi experiments, virgin females carrying the *Ubi-Gal4, ptub1□-Gal80^ts^* (*Ubi-Gal4, Gal80^ts^*) transposon were crossed to males carrying an UAS-RNAi transgene (TRiP) from the Tsinghua RNAi Center: THU02336.N (*CG18278* KD). The control flies were the offspring of the cross of the driver to UAS-mCherry RNAi VALIUM20 (Bloomington Stock Center # BL35785). Crosses with the *Ubi-Gal4, Gal80^ts^* driver were performed at 25°C for three days, then the progeny was left to develop at the non-permissive 18°C temperature. The hatched flies were kept at 29°C for five days prior to the experiment to allow Gal4-mediated transcription. All crosses involving flies without RNAi expression were performed at 25 °C. Unless stated otherwise, flies were five to seven day old.

To generate axenic flies, standard fly media was autoclaved. Antibiotics were added (Ampicillin 50ug/mL, Kanamycin 50ug/mL, Tetracyclin 50ug/mL, Erythromycin15ug/mL) when it cooled down to 50-60°C. The embryos were bleached then cultured on the sterilized media. The sterility of axenic flies (20 days old) was checked on LB, BHB, YPD, and MRS plates.

### Generation of CRISPR/Cas9-mediated null mutants

The *BaraA (CG18279)* and *CG18278* null mutants were generated using CRISPR/Cas9 technology based on the expression of gRNA transgenes that were then crossed to a transgenic line expressing a *pnos-Cas9* transgene. The 20bp-long gRNAs for the target genes were devised using web-based CRISPR Optimal Target Finder (http://targetfinder.flycrispr.neuro.brown.edu/). The plasmids carrying DNA sequences for the production of single strand gRNAs were constructed using standard methods. Briefly, the oligonucleotides were synthesized, denatured, and annealed to get double strand DNA before ligation into the expression vector, in which the gRNA coding sequences were transcribed under the control of the U6:3 promoter.

Plasmids carrying different gRNA targets were grouped by three or six for microinjection to obtain the gRNA transgenic fly lines, which were checked by sequencing. The gRNAs expressing plasmids were designed to be inserted on the 3^rd^ chromosome using *y^1^* M{vas-int.Dm}ZH-2A w*; M{3xP3-RFP.attP}ZH-86Fb (BL24749) flies. The gRNA flies were balanced before being crossed to flies carrying the *nosP*-*Cas9* transgene, to induce inheritable mutations. The primers used to generate the knock out mutants are shown in Table S1.

### Knock-in strategy

PCRs were done with the Q5 Hot-start 2× master mix (New England BioLabs, NEB), and cloning was performed using the Gibson Assembly 2× Master Mix (NEB) following the manufacturer’s instructions. The pCFD5 (U6:3-(t:: RNA^Cas9^)) plasmid vector was used. A cloning protocol to generate the pCFD5 plasmids encoding one to six tRNA-flanked sgRNAs was followed as described(45). The primers used to generate the pCFD5 vector containing the gRNAs are shown in Table S1. We used a pSK vector as donor plasmid with the homology arms flanking the mCherry: a fragment 1552bp upstream of *BaraA* had been amplified as a left arm; a fragment 1952bp downstream of *CG30059* as a right arm. Left arm + mCherry + right arm have been assembled (Gilson Assembly) and the resulting fragment ligated to Pst1-Spe1 double-digested pSK and checked by sequencing. The plasmid mixture containing the two plasmids at a ration pCFD5:pSK=3:1, was injected into recipient *y^1^* M{Act5C-Cas9.P.RFP-}ZH-2A *w^1118^* DNAlig4^169^ embryos.

### Overexpression strategy

Normal PCRs in first and second round were performed to amplify the ORF of BaraA or BaraA-derived peptides (DIM24, 12, 10, 13), constructing in pDONR221 with attP site (46). The primers used are shown in Table S1. BP recombination reaction was performed using with DH5α competent cells (Invitrogen); next, sequence-confirmed ORF entry clones were transferred to the destination vector pGW-HA.attB using a Gateway LR reaction (Gibson assembly). After validation by sequencing, the plasmids were injected in a pool into *y^1^* M{vas-int.Dm}ZH-2A w*; M{3xP3-RFP.attP}ZH-86Fb embryos and missing constructs were reinjected alone.

#### Pathogen infections

The bacterial strains used in this study include the Gram-negative bacterium *Pectinobacterium carotovorum carotovorum 15* (strain *Ecc15*, OD_600_=50) and the Gram-positive strains *Enterococcus faecalis (ATCC 19433)* (OD=0.1), *Micrococcus luteus* (OD=200) and *Staphylococcus albus* (OD=10), as well as *entV*^-^ and complemented *entV*^+^/*entV*^-^ and, which are derivatives of the wild-type *E. faecalis* OG1RF strain (OD=0.5) (kind gifts of Profs. Garsin and Lorenz, Houston, USA) (34). The fungal strains we used include filamentous fungi, *Aspergillus fumigatus* (5×10^7^ spores/mL, 250spores in 4.6nL)*, Metarhizium robertsii* (1×10^7^ spore/mL, natural infection 5×10^4^/mL), *DestruxinS1* mutant strain (1×10^7^ spore/mL), a kind gift from Prof. Wang, Shanghai, China (32). Besides, we used yeast as well, *Candida albicans* (pricked) and *Candida glabrata* (1×10^9^ yeasts/mL). The following media were used to grow the strains: Yeast extract-Peptone-Glucose Broth Agar (YPDA, *C. albicans* and *C. glabrata*) or Luria Broth (LB - all others) at 29°C (*Ecc15, M. luteus, C. albicans, C. glabrata*) or 37°C, *entV*^-^, *entV*^+^/*entV*^-^ *E. faecalis*, BHI medium, 37°C overnight, Rifampicin 100ug/ml. Spores of *M. robertsii* and *A. fumigatus* were grown on Potato Dextrose Agar (PDA) plates at 25°C or 29°C (*A. fumigatus*) for approximately one weeks or three weeks (*A. fumigatus)* until sporulation. We injected 4.6 nL of the suspension into each fly thorax using a Nanoject III (Drummond). Natural infections were initiated by shaking anesthetized flies in 5ml 0.01% tween-20 solution containing *M. robertsii* conidia at a concentration of 5×10^4^/mL. Infected flies were subsequently maintained at 29°C (*C. albicans, C. glabrata, A. fumigatus, M. robertsii*) or at 25°C (for all other pathogens, except for experiments with RNAi KD flies performed at 29°C). Flies were anesthetized with light CO2 for about three minutes during the injection procedure and were observed 3h after injection to confirm recovery from manipulations. Survival experiments were usually performed on three batches of 20 flies tested in parallel and pooled for statistical analysis using the Log-rank test.

### Pathogens Load Quantification

To characterize the dynamics of within-host microbial loads or BLUDs or FLUDs, live flies were taken at each time point post-injection for set point pathogen load or flies were infected with *E. faecalis* or *M. robertsii* and vials were monitored every 30 minutes for newly dead flies (PLUD). These flies were then individually homogenized with a bead in 100 μl PBS with 0.01% tween20 (PBST) or PBS. Homogenates were diluted serially in PBST (or PBS) and spread on LB (*E. faecalis*) or PDA (*M. robertsii*) plates for incubation at 37°C (*E. faecalis*) or 25°C *(M. robertsii)* overnight. Colonies were counted manually. Data were obtained from at least three independent experiments and pooled.

#### Gene Expression Quantitation

Total RNA was extracted from 5 adult flies collected at different time points after Toll activation with TRIzol (Invitrogen). cDNA was synthesized from 1000 ng total RNA using the SuperScript II Reverse Transcriptase kit (Transgene). Quantitative RT-PCR was performed on an iQ5 cycler (BioRad) using SYBR Green Supermix (Vazyme). Quantification of mRNA levels was calculated relative to levels of the ribosomal protein gene *rpl32*. See Table S2 for qPCR primer sequences. Primer couples displayed a similar efficiency, which allowed us to use the □□Ct method to analyze the data.

#### Survival tests

Survival tests were performed using 20-25 flies per vial in biological triplicates. Adult flies used for survival tests were 5–7-day old. For survival tests using RNAi-silencing genes, flies crossed at 25°C for 3 days for laying eggs then transferred to 18°C; after hatching, flies were kept for at least 5 days at 29°C prior to infections. Flies were counted every day. Each experiment shown is representative of at least two independent experiments.

#### Molecular mass fingerprints by MALDI MS

##### Sample preparation, data acquisition and processing

Each individual hemolymph sample was analyzed with the Bruker AutoFlex™ III based on Bruker Daltonics’ smartbeam laser technology. The molecular mass fingerprints (MFP) were acquired using a sandwich sample preparation on a MALDI MTP 384 polished ground steel plate (Bruker Daltonics Inc., Germany). Briefly, the hemolymph samples were 10-fold diluted in acidified water (0.1% trifluoroacetic acid - 0.1% TFA, Sigma Aldrich, France), 0.6μL was deposited on a thin layer of an air-dried saturated solution (0.6μL) of the matrix alpha-cyano-4-hydroxycinnamic Acid (4-HCCA, Sigma Aldrich, France) in pure acetone. Then 0.4 μL of a saturated solution of 4-HCCA prepared in 50% acetonitrile acidified with 0.1% TFA was mixed with the *Drosophila* hemolymph. Following co-crystallization of the hemolymph spots with the second matrix droplet and evaporation under mild vacuum, MALDI MS spectra were recorded in a linear positive mode and in an automatic data acquisition using FlexControl 4.0 software (Bruker Daltonics Inc.). The following instrument settings were used: the pseudo-molecular ions desorbed from the hemolymph were accelerated under 1.3kV, dynamic range of detection of 600 to 18,000 Da, between 50-60% of laser power, a global attenuator offset of 60% with 200Hz laser frequency, and 2,000 accumulated laser shots per hemolymph spectrum. The linear detector gain was setup at 1,906V with a suppression mass gate up to *m/z* 600 to prevent detector saturation by clusters of the 4-HCCA matrix. An external calibration of the mass spectrometer was performed using a standard mixture of peptides and proteins (Peptide Standard Calibration II and Protein Standard Calibration I, Bruker Daltonik) covering the dynamic range of analysis. All of the recorded spectra were processed with a baseline subtraction and spectral smoothing using FlexAnalysis 4.0 software (Bruker Daltonics Inc.).

#### Metabolite measurements

Female flies were first infected by 4.6nl *E. faecalis* or *M, robertsii* for 24 hours. Five flies per condition were used to measure the metabolite levels. Glycogen, glucose and triglycerides (TAGs) were measured using the Glycogen Assay Kit (700480), the Glucose Colorimetric Assay Kit (10009582), the Triglyceride Colorimetric Assay Kit (10010303 from Cayman Chemical), respectively. Standard curves were obtained according to the manufacturer’s instructions. The flies selected for each experiment had approximately the same body weight.

#### DestruxinA toxin injection

Destruxin A (MCE) was resuspended in high-quality grade DMSO and was diluted fivefold in PBS to a 1mM concentration. 23 nL of the solution or of control DMSO diluted in PBS at the same concentration was injected into flies using the Nanoject III microinjector (Drummond).

#### *Collection and preparation of* E. faecalis *supernatants*

Filter-sterilized supernatants phases were obtained from 10ml overnight cultures grown in LB medium that were collected by centrifugation at 4,000 rpm for 10min. The supernatants were sterilized by passing through a 0.2-μm-pore-size sterile syringe filter. The sterilized supernatants were centrifuged through a 15mL Amicon Centricon filter (Millipore) to separately collect the molecules larger or lower than 10kDa. 1.5mL Eppendorf tubes were used to collect the supernatant lower than 10kDa, which were vacuum freeze-dried for 24 hours. The powder was resuspended with H2O and thus concentrated 10 to 20-fold. The solution was filtered on 3kDa Amicon Centricon filter columns (Millipore) by centrifugation at 10000rpm for 30min. The nonfiltered fraction was then injected into flies with a volume optimized according to the batch (16 to 69 nL) and the same volume of buffer was used for the controls. All experiments were performed at least three times.

#### Statistics

Data are expressed as means ± SD. Data were analyzed by ANOVA (one-way) with Dunnett’s multiple comparisons test, with a significance threshold of P = 0.05. Log-rank tests were used to determine whether survival curves of female flies were significantly different from each other. Details are included in the legend of each figure. * p < 0.05; ** p < 0.01; *** p< 0.001; **** p<0.0001.

## Acknowledgments

We thank our colleagues at the Sino-French Hoffmann Institute for help and advice at various steps of this work. We are grateful to Dr. Wenhui Wang and Prof. Erjun Ling for providing antibodies, to Prof. Chengshu Wang for the *M. robertsii Destruxin*^-^ mutant strain, to Profs. Danielle Garsin and Michael Lorentz for the *entV* and *gelE* mutant strains (kindly sent by Melissa Cruz, Murray laboratory), to Dr. Lucas Waltzer for the PPO2-Gal4 stock, and the Bloomington and Tsinghua stock Centers for fly stocks. We would like to thank Sébastien Voisin from the BioPark for his help in mass spectrometry and Yongfang Zheng for expert technical help. We are indebted to the Guangzhou Drosophila Resource Center for the generation of CRISPR-Cas9 KO lines. We also thank Bruno Lemaitre and Mark Hanson for their cooperation and discussions on the *BaraA* gene.

