## Supplementary Information PDF for "*A Toll pathway effector protects* Drosophila *specifically from distinct toxins secreted by a fungus or a bacterium*"

**This PDF file includes:**

Figures S1 to S8
Tables S1-S2

A

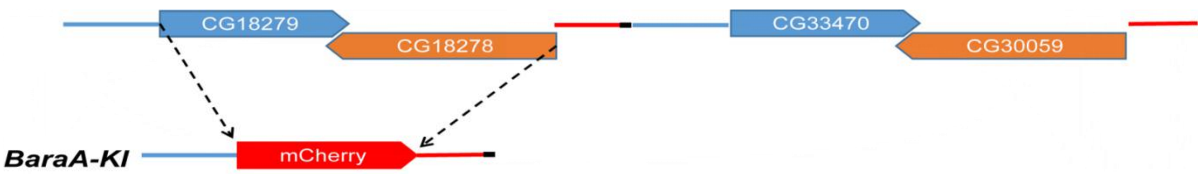


B

| | A5001 | yw | Canton-S | $W^{1118}$ | Ubi.Gal4. Gal80 | mCherry | WT | ΔBaraA-KO1 | ΔBaraA-KO2 | ΔBaraA-KI |
|---|---|---|---|---|---|---|---|---|---|---|
| 1 copy | ✔ | ✔ | ✔ | ✔ | ✔ | ✔ | ✔ | ✔ | ✔ | ✔ |
| 2 copies | | | | ✔ | | | | | | |

C

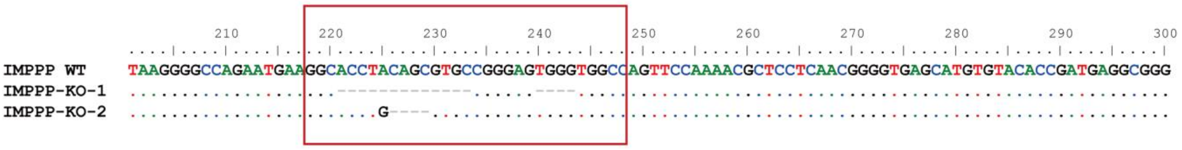


D

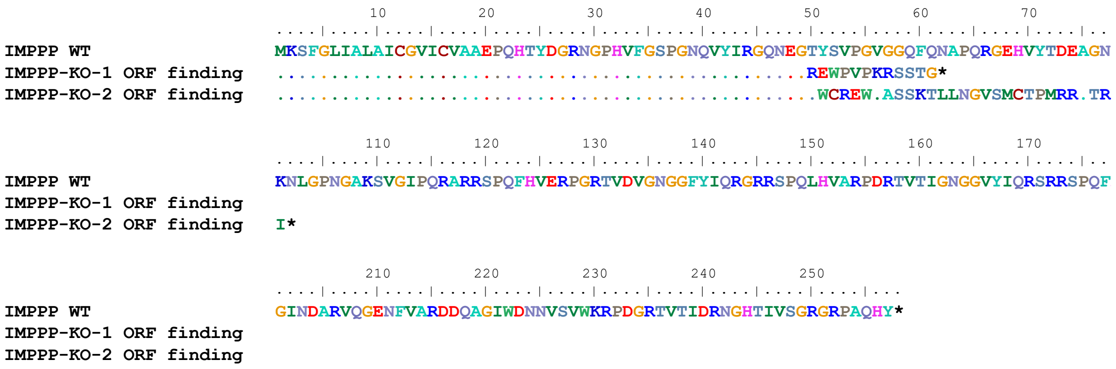


**Figure S1. Mutants affecting the *BaraA* locus.**

**(A)** Scheme of the tandem duplication of the *BaraA/CG18278(CG30059)* locus according to the *Drosophila* genome sequence; *CG18279* and *CG33470* (*BaraA*) on the one hand and *CG18278* and *CG30059* are perfectly duplicated, including 1172bp 5' to *CG18279* or *CG33470* start codon (shown as a blue line) and 774bp 5' to *CG18278* or *CG30059* start codon (shown as a red line). The black line represents the short unique region at the overlap of the duplicated loci. In the KI fly line, the two genes (*CG18279*, *CG18278*) were replaced by *mCherry* coding sequence after the START codon from *CG18279*. **(B)** Table recapitulating the tested strains and the presence of the duplication. The KI line was originally generated in a *yw* background with only one copy of the locus. **(C)** CRISPR Cas9 knock out mutants of *BaraA*: KO1 has a complex deletion pattern removing 17bp in total while the KO2 has a 4bp deletion and one point mutation. **(D)** The small deletions found in the KO lines lead to frame shift mutations which generate early stop codons.

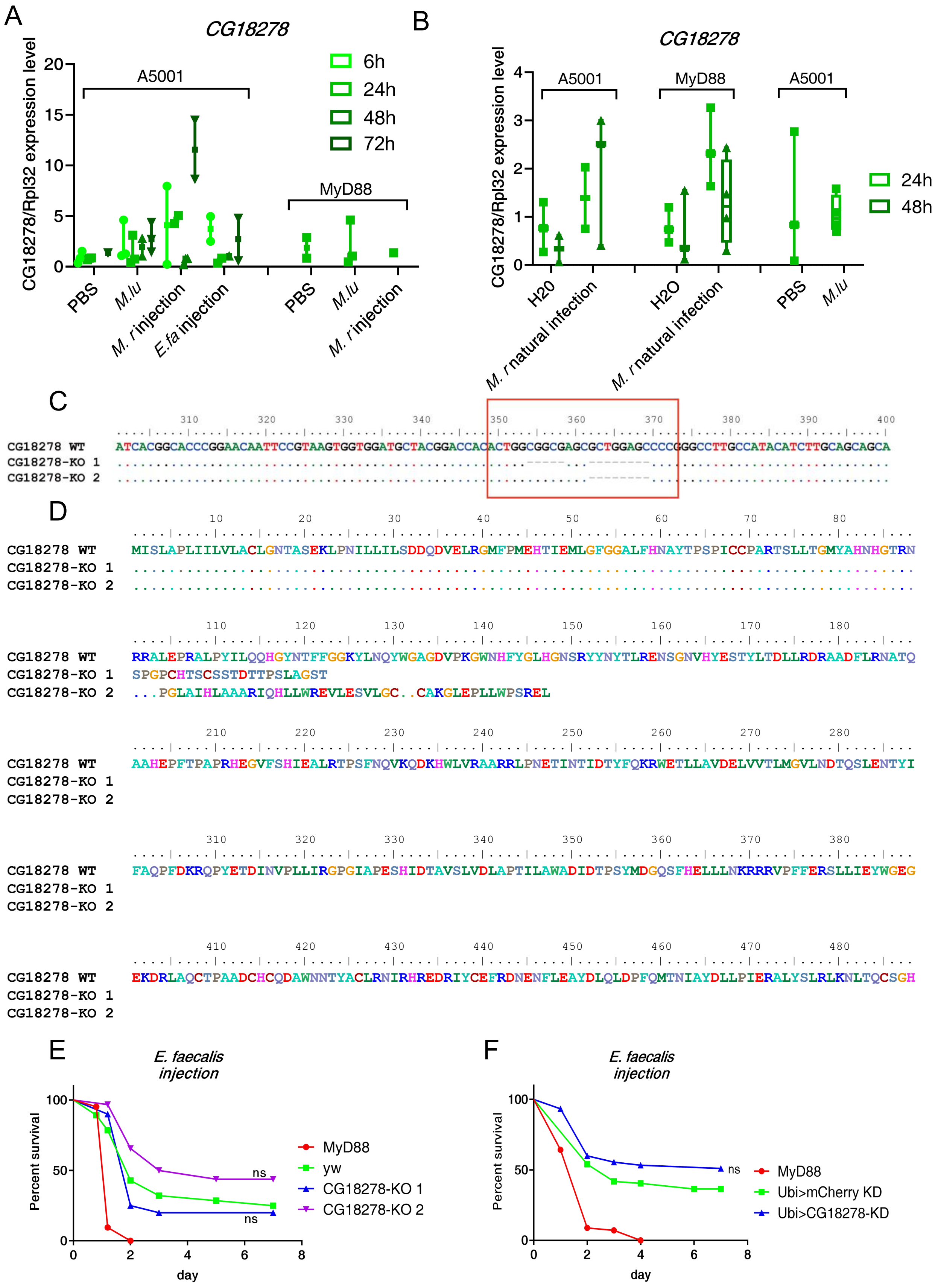
A
CG18278
CG18278/Rpl32 expression level
A5001
MyD88
6h
24h
48h
72h
PBS
M.lu
M. r injection
E.fa injection
PBS
M.lu
M. r injection
B
CG18278
CG18278/Rpl32 expression level
A5001
MyD88
A5001
24h
48h
H20
M. r natural infection
H2O
M. r natural infection
PBS
M.lu
C
CG18278 WT
CG18278-KO 1
CG18278-KO 2
ATCACGGCACCCGGAACAATTCCGTAAGTGGTGGATGCTACGGACCACACTGGCGGCGAGCGCTGGAGCCCCGGGCCTTGCCATACATCTTGCAGCAGCA
D
CG18278 WT
CG18278-KO 1
CG18278-KO 2
MISLAPLIILVLACLGNTASEKLPNILLILSDDQDVELRGMFPMEHTIEMLGFGGALFHNAYTPSPICCPARTSLLTGMYAHNHGTRN
RRALEPRALPYILQQHGYNTFFGGKYLNQYWGAGDVPKGWNHFYGLHGNSRYYNYTLRENSGNVHYESTYLTDLLRDRAADFLRNATQ
SPGPCHTSCSSTDTTPSLAGST
...PGLAIHLAAARIQHLLWREVLESVLGC..CAKGLEPLLWPSREL
AAHEPFTPAPRHEGVFSHIEALRTPSFNQVKQDKHWLVRAARRLPNETINTIDTYFQKRWETLLAVDELVVTLMGVLNDTQSLENTYI
FAQPFDKRQPYETDINVPLLIRGPGIAPESHIDTAVSLVDLAPTILAWADIDTPSYMDGQSFHELLLNKRRRVPFFERSLLIEYWGEG
EKDRLAQCTPAADCHCQDAWNNTYACLRNIRHREDRIYCEFRDNENFLEAYDLQLDPFQMTNIAYDLLPIERALYSLRLKNLTQCSGH
E
E. faecalis injection
Percent survival
day
MyD88
yw
CG18278-KO 1
CG18278-KO 2
ns
ns
F
E. faecalis injection
Percent survival
day
MyD88
Ubi>mCherry KD
Ubi>CG18278-KD
ns

**Figure S2. The *BaraA* neighboring gene *CG18278* is not involved in host defenses against *E. faecalis*.**

**(A)** Expression of the *CG18278* gene monitored by RTqPCR in wild-type $w^{A5001}$ and *MyD88* mutant flies at various time points after the injection of the indicated microbes; *M. lu: M. luteus*; *M. r: M. robertsii*; *E. fa: E. faecalis.* **(B)** Expression of the *CG18278* gene monitored by RTqPCR at 24 and 48 hours after a "natural" *M. robertsii* infection achieved by plunging the flies in a solution of conidia. Data were normalized to $w^{A5001}$ with *M. luteus* challenged after 24hours. Ct values from the RTqPCR for *CG18278* were in the 35-38 range while the Ct value for the Rpl32 were in the 18-20 range, which indicates that *CG18278* has low basal expression in fly. **(C, D)** Two lines of the CRISPR Cas9 mutant have been generated: *CG18278*-KO1 has a set of two small deletions removing altogether 13bp deletion whereas 8bp are deleted in the *CG18278* KO2 line. These deletions lead to frame shift mutations and early stop codons (D). **(E)** Nonisogenized *CG18278* KO mutants behave like the *yw* reference line when infected by *E. faecalis*. Two independent experiments have been performed and yielded similar results. **(F)** Flies in which *CG18278* is attenuated by RNAi KD driven by *Ubi*-Gal4 displayed a sensitivity to *E. faecalis* similar to that of the wild type. Two independent experiments have been performed.

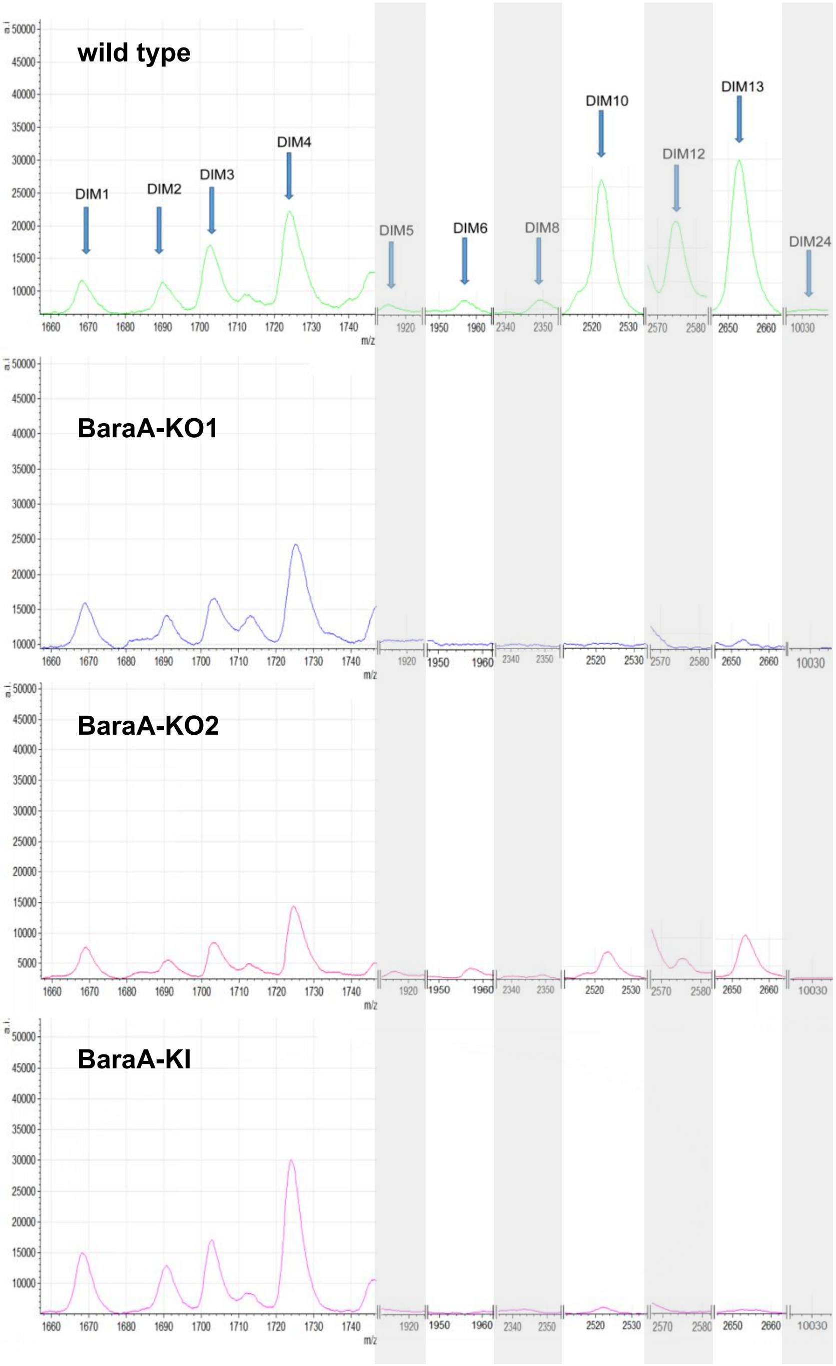


**Figure S3. Detection of *BaraA*-derived DIM peptides in wild-type and *BaraA* KD, KO, and KI mutants by mass-spectrometry analysis.**

The hemolymph was collected from single flies; four single flies were analyzed per genotype and yielded similar spectra by MALDI-TOF mass spectrometry. *BaraA* derived peptides were induced in wild type fly. Unexpectedly, in *BaraA*-KO2, DIM6, DIM10, DIM12, and DIM13 or unrelated peptides of similar molecular weight appear to be somewhat expressed. Unless otherwise specified, the notation *BaraA* KO refers to the KO1 line.

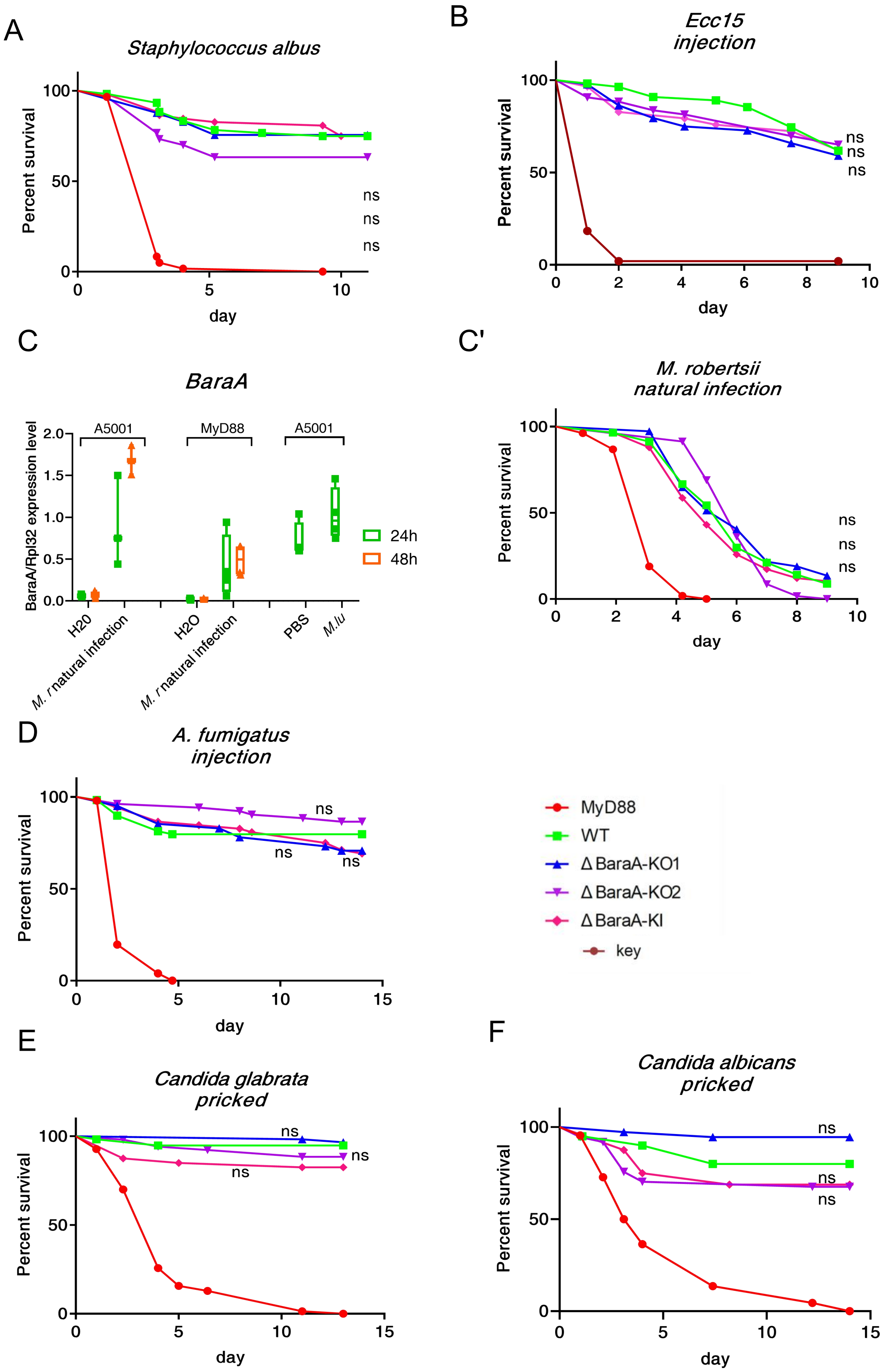

A
Staphylococcus albus
Percent survival
day
ns
ns
ns
B
Ecc15 injection
Percent survival
day
ns
ns
ns
C
BaraA
BaraA/Rpl32 expression level
A5001
MyD88
A5001
H20
M. r natural infection
H2O
M. r natural infection
PBS
M.lu
24h
48h
C'
M. robertsii natural infection
Percent survival
day
ns
ns
ns
D
A. fumigatus injection
Percent survival
day
ns
ns
ns
MyD88
WT
Δ BaraA-KO1
Δ BaraA-KO2
Δ BaraA-KI
key
E
Candida glabrata pricked
Percent survival
day
ns
ns
ns
F
Candida albicans pricked
Percent survival
day
ns
ns
ns

**Figure S4: *BaraA* mutants survive as well as wild-type (WT) flies to different types of infection.**

**(A, B; C'-E')** Survival experiments after the indicated infectious challenge in the septic injury model and natural infection model are presented and are representative of at least two independent experiments. The appropriate controls for the different microbes have been used: Gram-positive bacteria, fungi: *MyD88*, mutant of the Toll pathway; Gram-negative bacteria: *key*, mutant of the Immune deficiency pathway. None of the *BaraA* mutants displayed a reproducible susceptibility or resistance to infection (KO, KI). We used the log-rank test to determine the significance between wild-type and mutant survival curves. **(C)** Expression of *BaraA* steady-state transcripts during *M. robertsii* natural infection as monitored by RTqPCR.

A

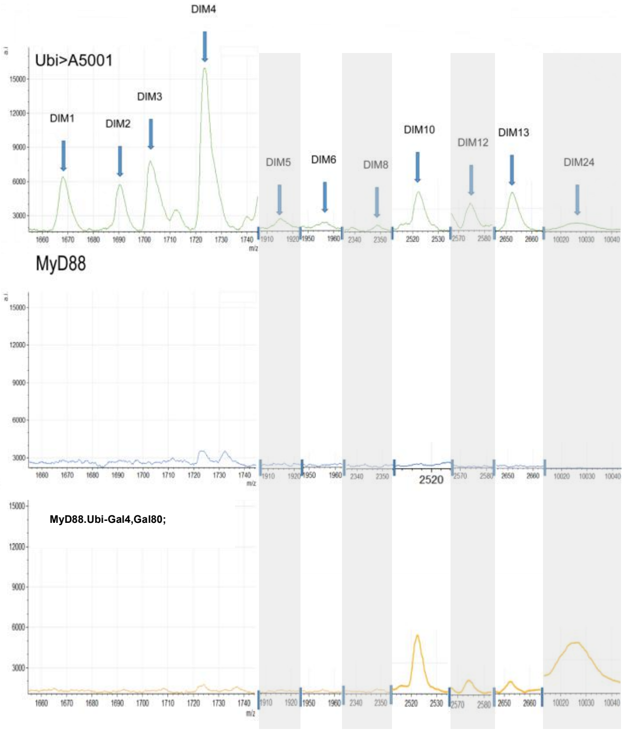


B

| | WT | MyD88 | BaraA-OE(MyD88) |
|---|---|---|---|
| DIM24 | 24819 | / | 134314 |
| DIM12 | 12949 | / | 4563 |
| DIM6 | 3047 | / | / |
| DIM10 | 24031 | / | 26381 |
| DIM13 | 18443 | / | 4211 |
| DIM8 | 2134 | / | / |
| DIM5 | 4332 | / | / |

C

| | | *E. faecalis* | *ECC15* | *M. robertsii* | *A. fumigatus* | *C. albicans* | *C. glabrata* |
|---|---|---|---|---|---|---|---|
| BaraA-HA or wt | OE in wt | 0 | 0 | 0 | 0 | 0 | 0 |
| | OE in MyD88 background | 0 | 0 | 0 | 0 | 0 | 0 |
| DIM24-HA or wt | OE in wt | 0 | 0 | 0 | 0 | 0 | 0 |
| | OE in MyD88 background | 0 | 0 | 0 | 0 | 0 | 0 |
| DIM12-HA or wt | OE in wt | 0 | 0 | 0 | 0 | 0 | 0 |
| | OE in MyD88 background | 0 | 0 | 0 | 0 | 0 | 0 |
| DIM10-HA or wt | OE in wt | 0 | 0 | 0 | 0 | 0 | 0 |
| | OE in MyD88 background | 0 | 0 | 0 | 0 | 0 | 0 |
| DIM13-HA or wt | OE in wt | 0 | 0 | 0 | 0 | 0 | 0 |
| | OE in MyD88 background | 0 | 0 | 0 | 0 | 0 | 0 |

D

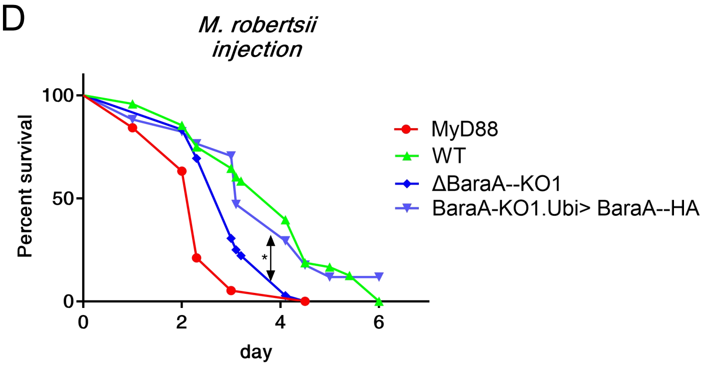


E

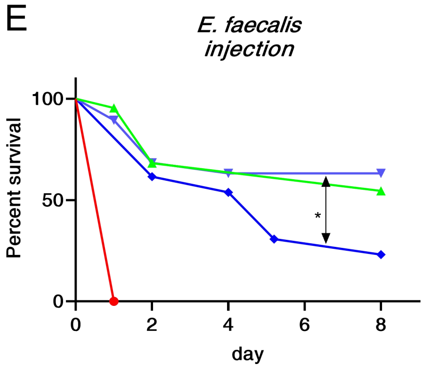

**Figure S5: *BaraA* overexpression in *MyD88* or WT background does not confer an enhanced protection against several pathogenic bacteria or fungi *in vivo*.**

**(A)** Mass-spectra of hemolymph from single flies was collected 24h after a *M. luteus* challenge for wild-type control flies, *MyD88* or flies overexpressing *BaraA* in a *MyD88* mutant background. In *MyD88* flies, only DIM4 (Daisho) was slightly expressed in contrast to the wild-type control in which DIM12, DIM10, DIM13 and DIM24 were detected, as well as other DIMs. Only the relevant parts of the spectra are shown. Quantification of the area under the curve shown in **(B)**. **(C)** We generated transgenes allowing the expression of *BaraA* or sequences encoding specific DIM domains derived from BaraA (DIM 22 excepted) under the control of UAS enhancer sequences and expressed them in WT or *MyD88* background: no altered survival to the indicated pathogens was observed, "0" means no phenotype. **(D, E)** Rescue of the nonisogenic *BaraA* KO1 mutant with a *BaraA*-HA (full BaraA coding sequences) expressed under the control of a p*Ubi*-Gal4$^{ts}$ driver (cross performed at 18°C and induced at the adult stage at 29°C) after *M. robertsii* (D) or *E. faecalis* injection (E). Three independent experiments with each pathogen have been performed. *$p < 0.05$.

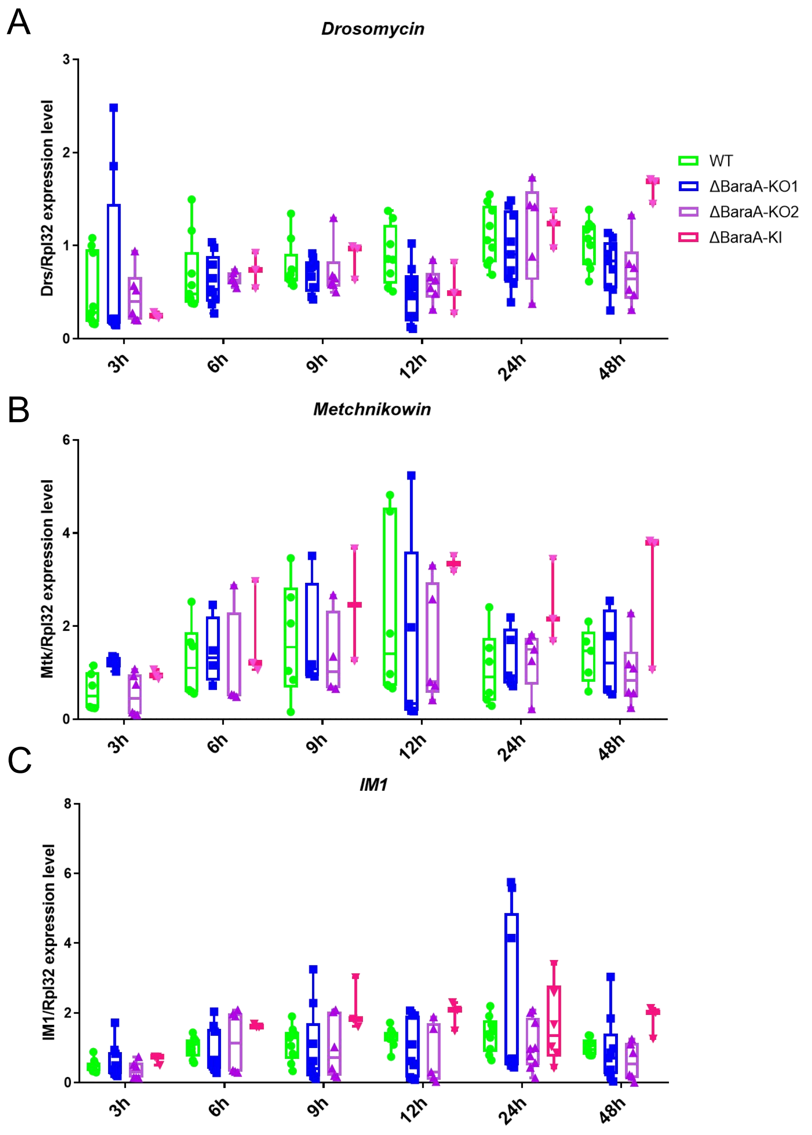


**Figure S6. Toll-mediated activation of some of its target effector genes is not altered in *BaraA* KO or KI flies.**

Steady-state transcript levels of *D. melanogaster* Toll pathway-regulated genes were measured by quantitative RT-PCR at different time points after a *E. faecalis* challenge: *Drosomycin* (A), *Metchnikowin* (B), and *DIM1=BomS1*(C). These experiments are representative of three independent experiments. Gene expression was normalized against *rpl32* gene expression and the results are normalized to the expression at 48h measured with WT. No significant difference between WT and isogenic *BaraA* mutants was detected. The Kruskall-Wallis multiple comparisons test has been used.

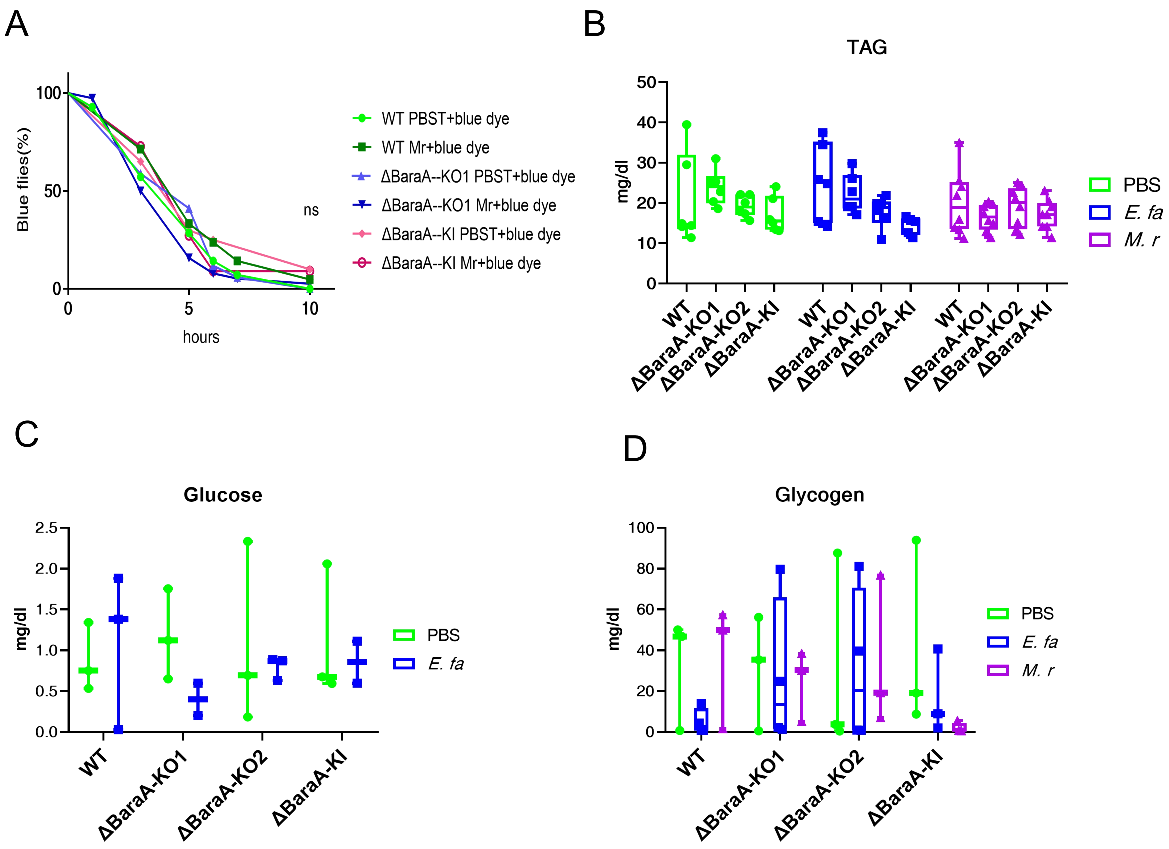


**Figure S7. *BaraA* does not appear to be involved in the function of Malpighian tubules nor in the management of metabolic stores during infection.**

**(A)** FRUMS assay: the flies were first injected with 50 *M. robertsii* conidia; 4.6nl blue dye was injected into flies 24hpi and the clearing of the dye from the abdomen was monitored the following hours. Quantification of FRUMS assay: two batches of twenty flies were monitored each hour. **(B-D)** Flies were injected either with PBS, *E. faecalis* or *M. robertsii*. 24 hours post infection, batches of five flies were homogenized in buffer, then centrifuged and the supernatants were used to measure the metabolic stores: triacylglycerides (TAG, B), glucose (C), and glycogen (D).

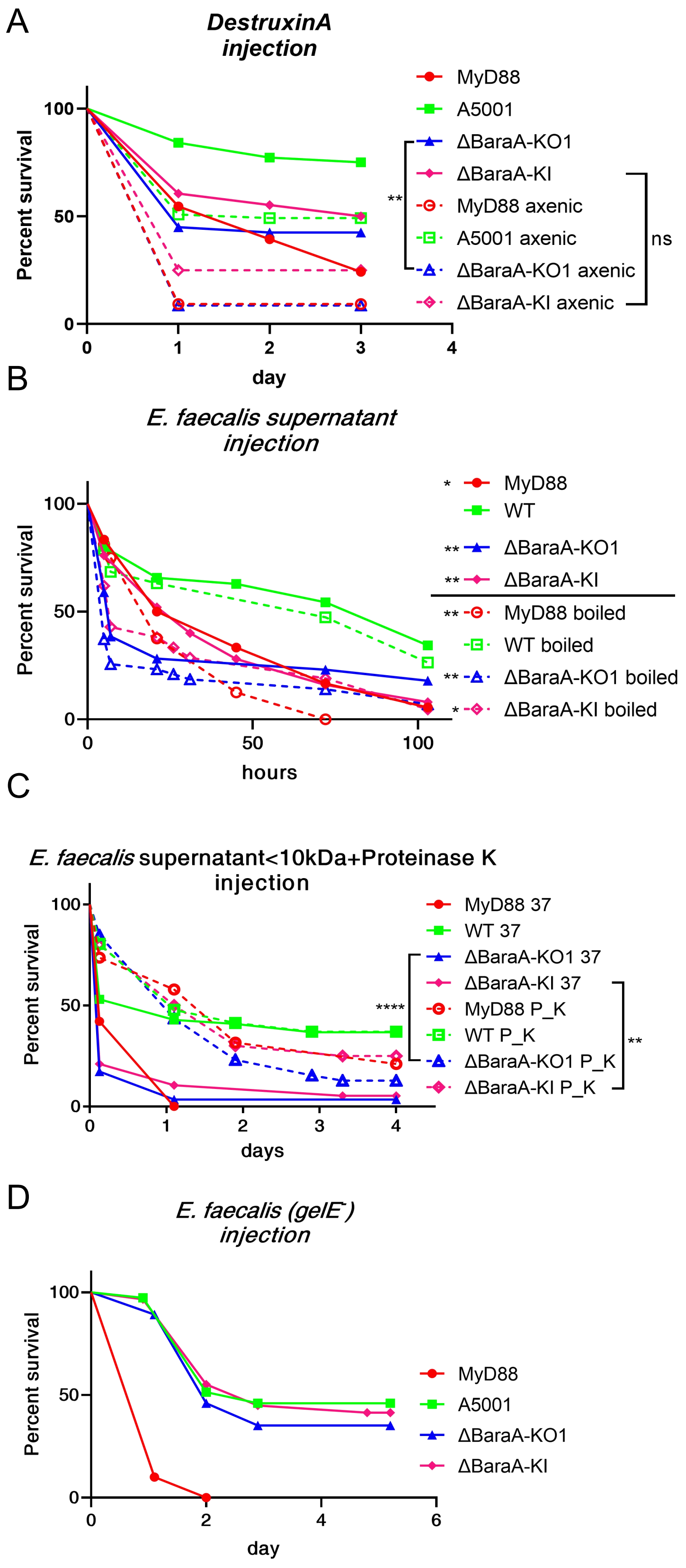
A
DestruxinA injection
Percent survival
day
MyD88
A5001
ΔBaraA-KO1
ΔBaraA-KI
MyD88 axenic
A5001 axenic
ΔBaraA-KO1 axenic
ΔBaraA-KI axenic
**
ns
B
E. faecalis supernatant injection
Percent survival
hours
MyD88
WT
ΔBaraA-KO1
ΔBaraA-KI
MyD88 boiled
WT boiled
ΔBaraA-KO1 boiled
ΔBaraA-KI boiled
C
E. faecalis supernatant<10kDa+Proteinase K injection
Percent survival
days
MyD88 37
WT 37
ΔBaraA-KO1 37
ΔBaraA-KI 37
MyD88 P_K
WT P_K
ΔBaraA-KO1 P_K
ΔBaraA-KI P_K
****
**
D
E. faecalis (gelE⁻) injection
Percent survival
day
MyD88
A5001
ΔBaraA-KO1
ΔBaraA-KI

**Figure S8. Further characterization of toxins secreted by *M. robertsii* or *E. faecalis***

**(A)** 4.6nL of 8mM DestruxinA were injected into axenic (dashed lines) or conventionally-raised flies. DestruxinA-treated *BaraA*-KO1 showed significant difference from control axenic *BaraA*-KO1, $^{**}p<0.01$ whereas only a trend was observed with the *BaraA*-KI line, suggesting that *BaraA* and *MyD88* flies do not succumb to an opportunistic infection. **(B)** Supernatant from *E. faecalis* was boiled at 95°C for 5min. Flies were injected with 23nl of the boiled (dashed lines) or untreated supernatant. For both conditions, injected mutant flies were significantly more susceptible than control WT flies. For each condition (above or below the line in the caption), mutant flies are compared to wild-type flies submitted to the same challenge for statistical analysis. * $p<0.05$; ** $p< 0.01$. **(C)** Supernatant was incubated with 100ug/ml Proteinase K or with PBS (the same volume as Proteinase K) at 37°C for 18 hours. 23nl of supernatant was injected. *BaraA*-KO1 with Proteinase K treated supernatant (dashed lines) died slower than the flies injected with untreated supernatant. The same result was observed for *BaraA*-KI. ** $p< 0.01$, **** $p< 0.0001$. (B, C) three independent experiments have been performed. **(D)** 4.6nL of 0.5 OD of the *GelE- E. faecalis* mutant strain was injected. No statistically significant difference was observed between wild type and *BaraA* mutant flies in four out of five experiments.

**Table S1: Primers used for cloning**

| | |
|---|---|
| BaraA Fw | AAAAAGCAGGCTTCAACATGAAATCGTTTGGATTGATTGC |
| BaraA Rv | AGAAAGCTGGGTCTTAAACTTTTTGGAGGCATATGA |
| BaraA Rv-HA | AGAAAGCTGGGTCAACTTTTTGGAGGCATATGA |
| DIM24 Fw | AAAAAGCAGGCTTCAACATGCAACACACCTACGACGGTC |
| DIM24 Rv | AGAAAGCTGGGTCTTAGGCACGTTGTGGGATTCCC |
| DIM24 Rv-HA | AGAAAGCTGGGTCGGCACGTTGTGGGATTCCC |
| DIM12 Fw | AAAAAGCAGGCTTCAACATGCAGTTTCACGTAGAGCGTC |
| DIM12 Rv | AGAAAGCTGGGTCTTAGCCACGCTGAATGTAGAATC |
| DIM12 Rv-HA | AGAAAGCTGGGTCGCCACGCTGAATGTAGAATC |
| DIM10 Fw | AAAAAGCAGGCTTCAACATGCAGCTTCATGTGGCACGCC |
| DIM10 Rv | AGAAAGCTGGGTCTTAACTGCGTTGAATATAGACGCC |
| DIM10 Rv-HA | AGAAAGCTGGGTCACTGCGTTGAATATAGACGCC |
| DIM13 Fw | AAAAAGCAGGCTTCAACATGCAGTTCCATGTGGAGCGAC |
| DIM13 Rv | AGAAAGCTGGGTCTTAGAAACGTTGAGCGGAGAAGC |
| DIM13 Rv-HA | AGAAAGCTGGGTCGAAACGTTGAGCGGAGAAGC |
| 2nd-F | GGGGACAAGTTTGTACAAAAAAGCAGGCT |
| 2nd-R | GGGGACCACTTTGTACAAGAAAGCTGGGT |
| BaraA gRNA-F(KI) | GCGGCCCGGGTTCGATTCCCGGCCGATGCAATCTGTGGC GTTATCTGCGTGTTTTAGAGCTAGAAATAGCAAG |
| BaraA gRNA-R(KI) | ATTTTAACTTGCTATTTCTAGCTCTAAAACCTGCCTTTTACA ACACTGCATGCACCAGCCGGGAATCGAACCC |
| BaraA gRNA | ACCCACTCCCGGCACGCTGT |
| CG18278 gRNA | ACCACACTGGCGGCGAGCGC |

**Table S2: Primers used for RTqPCR**

| | |
|---|---|
| RpL32 Fw | GCTAAGCTGTCGCACAAATG |
| RpL32 Rv | GTTCGATCCGTAACCGATGT |
| Drosomycin Fw | TACTTGTTCGCCCTCTTCG |
| Drosomycin Rv | GAGCGTCCCTCCTCCTTGC |
| IM1 Fw | CAATGCTGTTCCACTGTCGC |
| IM1 Rv | CGTGGACATTGCACACCCTG |
| Metchnikowin Fw | CGTCACCAGGGACCCATTT |
| Metchnikowin Rv | CCGGTCTTGGTTGGTTAGGA |
| BaraA Fw | GGTGAGCATGTGTACACCGA |
| BaraA Rv | GGCGGAAAAATTGGGACCAC |
| CG18278 Fw | GCCCATGAACCCTTCACTCC |
| CG18278 Rv | CACCAACCAGTGCTTATCCTGC |
